# Reducing astrocyte calcium in the nucleus accumbens core increases reward valuation

**DOI:** 10.1101/2025.10.20.683571

**Authors:** Jonathan W VanRyzin, Gabrielle A Laraia, Utsav Gyawali, Morgan H James, Kathryn J Reissner

**Affiliations:** Department of Psychology and Neuroscience, University of North Carolina at Chapel Hill, Chapel Hill, NC, USA; Department of Psychiatry, Robert Wood Johnson Medical School, Rutgers University, Piscataway, NJ, USA; Rutgers Addiction Research Center, Brain Health Institute, Rutgers Health, Piscataway, NJ, USA; School of Psychology, Faculty of Science, University of Sydney, Sydney, New South Wales, Australia; Brain and Mind Centre, University of Sydney, Sydney, New South Wales, Australia

## Abstract

Astrocyte dysfunction within the nucleus accumbens (NAc) has been increasingly implicated in maladaptive reward processing and the development of addiction-like behaviors. In this study, we investigated how impairing astrocyte calcium signaling in the NAc core influences consummatory reward behaviors and reward valuation. Using a viral approach to express the human plasma membrane calcium ATPase (hPMCA) in NAc core astrocytes in male rats, we selectively reduced intracellular astrocyte calcium dynamics. Blunting astrocyte calcium signaling led to increased self-administration of both sucrose and cocaine on a low effort fixed-ratio schedule of reinforcement. Behavioral economic analysis revealed enhanced reward motivation and valuation in hPMCA-expressing rats as compared to controls. Notably, reduction of astrocyte calcium signaling did not alter cocaine-induced locomotor sensitization, indicating a dissociation between astrocytic regulation of reward valuation and other cocaine-related behaviors. Collectively, these results identify astrocyte calcium dynamics in the NAc core as a key constraint on motivational drive and reward valuation.

## INTRODUCTION

The nucleus accumbens (NAc) is a central hub in the brain’s reward circuitry, involved in processing reward-related information, motivation, and guiding subsequent action selection (Mogenson et al., 1980). The NAc core subregion, in particular, processes reward-related information to evaluate the valuation and salience of reward and facilitate the acquisition of self-administration, reinstatement, and seeking behaviors in preclinical rodent models of drug use (Lobo et al., 2010; Stuber et al., 2011; Calipari et al., 2016; Scofield et al., 2016; Soares-Cunha et al., 2016; Kutlu et al., 2021). While these models have been extensively employed to inform neuronal mechanisms of reward processing, the contributions of non-neuronal cells remain poorly understood.

Recent evidence indicates that astrocytes play critical roles in shaping NAc circuit function and reward-seeking behavior (Scofield et al., 2016; Kruyer & Scofield, 2021; Harder et al., 2024). Astrocytes dynamically influence neural circuit function through neurotransmitter uptake, release of gliotransmitters, and modulation of synaptic strength and plasticity (Dallérac et al., 2018; Nagai et al., 2021; Lyon & Allen, 2022). In models of substance use disorders, NAc core astrocytes undergo structural and functional changes, leading to atrophy and withdrawal from synapses, and impaired glutamate homeostasis (Kalivas, 2009; Wang et al., 2022). These changes are thought to contribute to the reinstatement of drug seeking and heightened vulnerability to relapse, highlighting the importance of astrocytes in maintaining homeostasis in reward circuits and the regulation of motivated behaviors.

The mechanisms by which astrocytes regulate neural circuits largely depend upon intracellular calcium signaling. Astrocytic calcium transients can be triggered by local release of neurotransmitters, such as glutamate and dopamine, leading to downstream release of gliotransmitters and modulation of neural activity (Papouin et al., 2017; Goenaga et al., 2023). In this way, calcium signaling enables astrocytes to sense and respond to local neural activity, and through interconnected astrocyte networks, link changes in synaptic dynamics to circuit-level regulation (Nagai et al., 2021). To causally investigate how astrocyte calcium dynamics contribute to reward-related behavior, recent studies have used AAV-mediated expression of the human plasma membrane calcium ATPase (hPMCA2w/b; “hPMCA”) to manipulate astrocyte calcium *in vivo*, which reduces basal calcium and decreases the amplitude and duration of spontaneous and evoked calcium responses in astrocytes (Yu et al., 2018). These studies show that hPMCA expression in dorsal stratum astrocytes increases cue-induced reinstatement of cocaine seeking (Tavakoli et al., 2024), while expression in cortical astrocytes reduces ethanol intake (Erickson et al., 2020) and prevents ethanol-induced changes in neuron firing rates (Kastner-Blascyzk et al., 2025).

Given these observations, we sought to determine how disrupting astrocyte calcium signaling in the NAc core alters consummatory behaviors for both natural and drug reinforcers in rats. We used AAV-mediated expression of hPMCA selectively in NAc core astrocytes to reduce intracellular calcium signaling and assessed consumption of sucrose and cocaine on a low effort fixed-ratio schedule, as well as behavioral economic parameters of valuation and motivation. We show that hPMCA expression in astrocytes increases self-administration of both sucrose and cocaine, accompanied by increased measures of demand, including higher essential value and maximum price paid, as well as lower demand elasticity. In contrast, hPMCA expression did not affect cocaine-induced locomotor sensitization, indicating that astrocytic calcium signaling specifically constrains reward consumption and valuation without broadly impairing drug-related plasticity. Together, these findings highlight astrocyte function in the NAc core as a key constraint on motivational drive and valuation of both natural and drug reinforcers.

## MATERIALS AND METHODS

### Animals

Adult male Sprague-Dawley rats (225-250 g; Envigo) were single-housed in an AAALAC-accredited animal facility at the University of North Carolina at Chapel Hill (UNC) on a 12:12 h reverse light/dark cycle with ad libitum food and water. Upon arrival, rats were allowed to acclimate for at least 1 week prior to the start of experimentation. All care and procedures were performed in accordance with the National Institutes of Health *Guide for the Care and Use of Laboratory Animals* and were approved by the Institutional Animal Care and Use Committee at UNC.

### Intracranial Viral Infusion

Rats were anesthetized with ketamine (100 mg/kg) and xylazine (7 mg/kg) and received bilateral stereotaxic infusion of either AAV5-GfaABC1D-mCherry-hPMCA2w/b (1 µl/hemisphere; 1.0 x 10^13^ viral particles/ml; Addgene #111568) or AAV5-GfaABC1D-Lck-mCherry (1 µl/hemisphere; 6.0 x 10^12^ viral particles/ml; plasmid was a generous gift from Joshua Jackson at Drexel University) directly into the NAc core (+1.5 AP, ±2.6 ML, -7.2 DV, 6° angle) using a 24G syringe (Hamilton) at a rate of 100 nl/min. Viruses were allowed to diffuse for an additional 10 min following infusion. Rats were allowed to recover for at least 2 weeks before behavioral testing to ensure proper viral expression.

### Intravenous catheter surgery

Rats were anesthetized with ketamine (100 mg/kg) and xylazine (7 mg/kg) and implanted with indwelling silastic catheters in the jugular vein as previously described (Testen et al., 2025). Post surgery, and each day subsequently, catheters were flushed with gentamycin (0.1 ml, 5 mg/ml) and heparin (0.1 ml, 100 U/ml) to maintain patency. Animals were allowed to recover for at least 5 days before behavioral testing. Catheter patency was verified prior to behavioral testing with a subthreshold dose of propofol (0.05 ml, 10 mg/ml).

### Food Training

Rats were food trained to lever press on a fixed-ratio 1 (FR1) reinforcement schedule for a single food pellet (45 mg, BioServe) for at least 2 h and until the number of active lever presses was >75 with a ratio of active:inactive lever presses of 2:1. Rats were kept on food restriction (∼20 g/day) for the duration of the experiment.

### Sucrose self-administration

Prior to sucrose self-administration, rats were food trained to lever press on an FR1 reinforcement schedule as described above. Rats were then tested for FR1 self-administration of a sucrose pellet (45 mg, Test Diet) for 2 h/day for 10 days. Testing occurred in operant chambers in sound-attenuating boxes controlled by Med-PC IV software (Med Associates). During each session, the house light was illuminated and rats were presented with an active and inactive lever. Successful active lever presses resulted in the delivery of one sucrose pellet which was paired with a white cue light and tone (70 dB, 2.5 kHz) for 5 s followed by a 20 s timeout period, during which further active lever presses failed to deliver sucrose pellets.

### Cocaine self-administration

Cocaine HCl (National Institute of Drug Abuse) was dissolved in sterile saline at a concentration of 5 mg/ml. Prior to cocaine self-administration, rats were food trained to lever press on an FR1 reinforcement schedule as described. Rats were then tested for FR1 self-administration of intravenous (i.v.) cocaine for 6 h/day for 10 days. Testing occurred in the same operant chambers as described above. Successful active lever presses resulted in an iv infusion of cocaine (0.75 mg/kg/infusion) and were followed by a 20 s timeout period, during which further active lever presses failed to deliver iv cocaine infusions.

### Sucrose behavioral economics

Rats were trained to lever press for sucrose pellets on an FR1 reinforcement schedule for 2 h/day as described above. After FR1 training, rats were tested daily on a 2 h between-session behavioral economics protocol as previously described (Heinsbroek et al., 2021), with increasing FR requirements each day as follows: FR1, FR3, FR5, FR10, FR18, FR32, FR56, FR100. Testing continued until each rat failed to receive one sucrose pellet.

### Cocaine behavioral economics

Rats were trained to lever press for i.v. cocaine (2.73 mg/ml) on an FR1 reinforcement schedule for 6 h/day as described above. After FR1 training, rats were tested on a within-session behavioral economics procedure, as described previously (Bentzley et al., 2012). Each session lasted 110 min and the dose of cocaine per infusion decreased in a quarter logarithmic scale across successive 10 min bins as follows: 383.5, 215.6, 121.3, 68.2, 38.3, 21.6, 12.1, 6.8, 3.8, 2.2, 1.2 μg cocaine per infusion. The amount of cocaine per infusion was determined by the duration of syringe pump activation following each lever press, which decreased with each succeeding 10 min bin. Rats were tested daily for 3 days and data were derived from the average across testing days.

### Economic demand curve fitting

Exponential demand curves were fit to each animal’s behavioral data using the exponential demand equation (Hursh & Silberberg, 2008). From the demand curves, the following parameters were calculated: α (demand elasticity), Q_0_ (theoretical consumption at zero cost), P_max_ (maximum effort for unit reward), and essential value (reinforcing efficacy of a reward).

### Behavioral sensitization of locomotor activity

Behavioral sensitization of locomotor activity was performed as previously described (Pierce et al., 1996; Boudreau & Wolf, 2005). Locomotor activity testing occurred in open arenas (17 x 17 x 12 in) equipped with infrared beams to allow for animal tracking (Med Associates). Arenas were housed in sound-attenuating chambers and controlled by Activity Monitor software (Med Associates). On the first day of behavioral testing, rats were placed into the center of the arena and locomotor activity was measured for 30 min to establish baseline activity. Rats were then injected with cocaine (15 mg/kg; i.p.), placed back into the arena, and locomotor activity was measured for an additional 2 h. For the next 5 days, rats were injected with cocaine (20 mg/kg; i.p.) in their home cage. On the last day of testing (day 7), rats were placed into the center of the testing arena to measure locomotor activity for 30 min, then injected with cocaine (15 mg/kg; i.p.) and placed back into the arena to measure locomotor activity for an additional 2 h.

### Immunohistochemistry

Rats were anesthetized with sodium pentobarbital, then transcardially perfused with phosphate buffered saline (PBS; 0.1M, pH 7.4) followed by 4% paraformaldehyde (PFA; 4% in PBS, pH 7.2). Brains were postfixed in 4% PFA for 24 h at 4°C, then kept in 30% sucrose solution at 4°C until fully submerged. Coronal sections of the nucleus accumbens were cut using a cryostat (Leica) at a thickness of 45 µm and stored in cryoprotectant solution at -20°C until processing.

Tissue sections were mounted onto slides and washed with PBS, then blocked with 5% bovine serum albumin (BSA) in PBS + 0.4% Triton X-100 (PBS-T) for 1 h. Slides were incubated in primary antibody solution (2.5% BSA in PBS-T) overnight at RT. The following day, slides were washed with PBS, incubated in secondary antibody solution (2.5% BSA in PBS-T) for 2 h, stained with Hoescht 33342 (1:2000; Invitrogen), and coverslipped with ProLong Diamond Antifade Mountant (Thermo Fisher Scientific). The following primary antibodies were used in these studies: chicken anti-mCherry (Abcam; #AB205402), rabbit anti-GFAP (Dako; #z0334) mouse anti-NeuN (EMD Millipore; #MAB377). The following secondary antibodies were used in these studies: Alexa Fluor (all 1:500; Thermo Fisher Scientific) donkey anti-488, 594, and 637.

### Confocal microscopy

Confocal fluorescent images were acquired using a Zeiss LSM 800 equipped with 405, 488, 561, and 640 lasers and a 20x (0.8 NA) and 63x (1.4 NA) oil-immersion objective. Images were acquired as z stacks using 2.5µm z-step for 20x images and 0.5 µm z-step for 63x images. All images were deconvolved using AutoQuant software (version X3.0.4; MediaCybernetics) and processed to correct for background in Imaris software (version 9.9; Bitplane).

### Cell Quantification

Quantification of mCherry-hPMCA2w/b expression with NeuN and GFAP was performed using Imaris software. At least two fields of view were imaged using a 20x objective with a 2x2 tile scan across two sections for each animal, and the number of mCherry+/NeuN+ and mCherry+/GFAP+ cells were quantified. Cells were included in the analysis if a distinct soma could be identified for each cell.

### Statistical analysis

Statistical analyses were performed using GraphPad Prism (version 10). Details of specific statistical analyses can be found in figure legends and in the test (e.g. tests used, exact n, p value). Self-administration data were analyzed using two-way repeated measures (RM) analysis of variance (ANOVA). Comparisons between two experimental groups were performed using a two-tailed t test for normally distributed data or Mann-Whitney U test for non-normally distributed data. The assumption of normality was assessed using Shapiro-Wilk test.

## RESULTS

### Expression of hPMCA in nucleus accumbens core astrocytes increases self-administration of sucrose and cocaine

To determine how impairing astrocyte calcium signaling within the NAc core would impact consummatory reward-related behaviors, we expressed astrocyte-specific hPMCA, which encodes a cell membrane calcium exporter, to extrude intracellular calcium. This construct has been previously shown to reduce basal calcium, as well as the amplitude and duration of spontaneous and evoked calcium responses in astrocytes (Yu et al., 2018). We bilaterally injected AAV5-GfaABC1D-mCherry-hPMCA2w/b or AAV5-GfaABC1D-Lck-mCherry as a control into the NAc core prior to behavioral analysis (Figure 1A). After 3 weeks of expression, we found that hPMCA expression was largely restricted to the NAc core (Figure 1B). Colocalization analysis with the astrocyte marker, GFAP and neuron marker, NeuN revealed that the vast majority (94.95% ± 1.71%) of mCherry+ cells co-labeled with GFAP, whereas far fewer (5.05% ± 1.71%) mCherry+ cells co-labeled with NeuN (Figure 1C, 1D).

**Figure 1.**
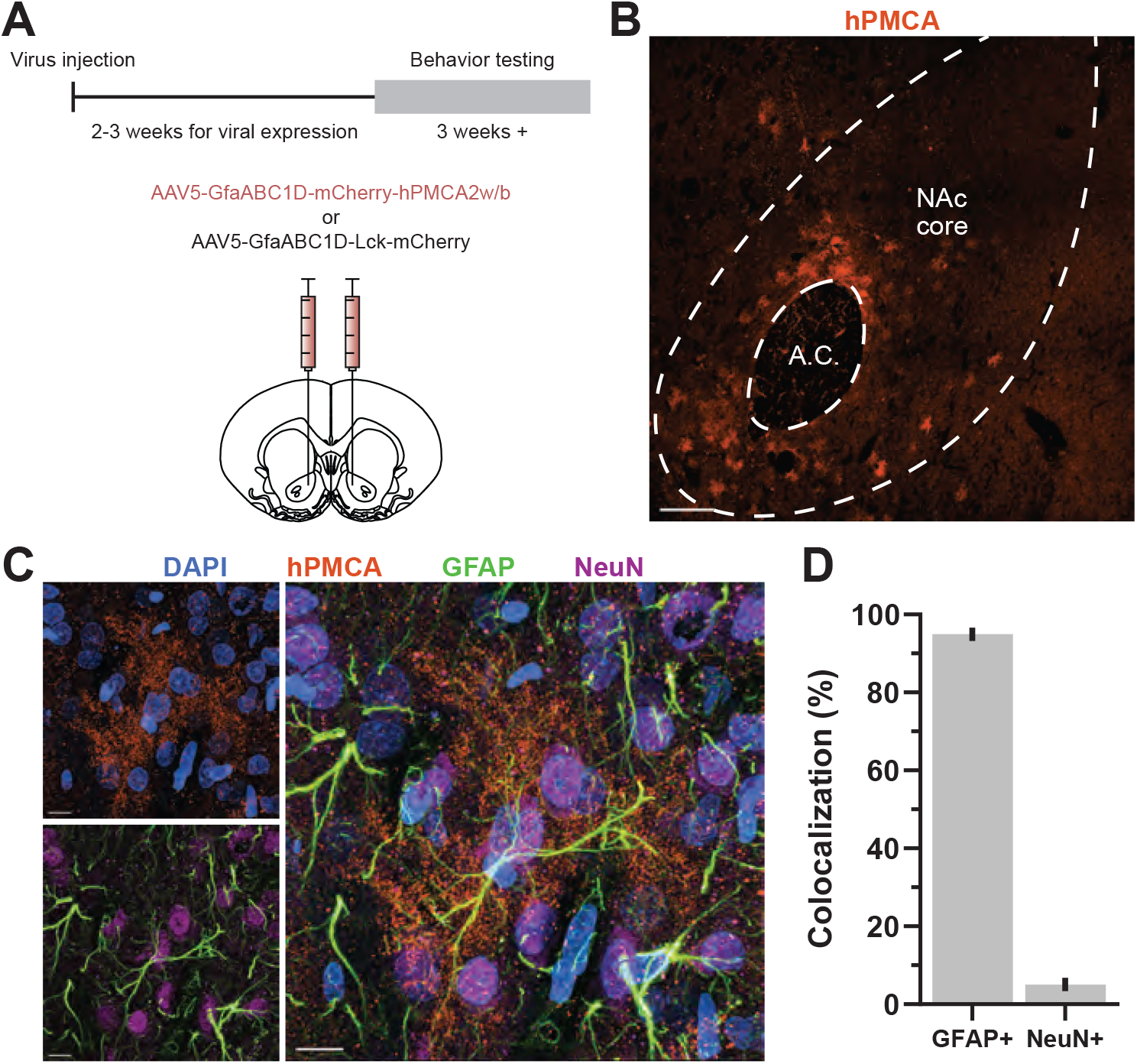
AAV5-GfaABC1D-mCherry-hPMCA2w/b expression in the NAc core is highly specific to astrocytes. **A)** Experimental timeline. AAV5-GfaABC1D-mCherry-hPMCA2w/b was injected into the NAc core 3-4 weeks prior to the start of self-administration or behavioral economic testing. **B)** Viral expression of hPMCA was highly localized to the NAc core. Scale bar = 200 µm. **C)** Representative high magnification image of DAPI and hPMCA expression showing classic astrocyte morphology (top left), the astrocyte- and neuron-specific markers, GFAP and NeuN, respectively (bottom left), and the resulting merged image (right). Scale bar = 10 µm. **D)** Quantification of hPMCA+ cells with either GFAP or NeuN shows highly specific expression restricted to astrocytes. n = 6 rats. Bars represent the mean ± SEM. NAc, nucleus accumbens.

We then tested rats for 2 h/day across 10 days of a sucrose self-administration paradigm with an FR1 reinforcement schedule to determine whether hPMCA expression would alter the consumption of a palatable food reward. Compared to mCherry controls, hPMCA rats obtained more sucrose pellets across test days (Figure 2A), with a corresponding increase in the number of active lever presses, but not inactive lever presses (Figure 2B). Importantly, both hPMCA and mCherry rats obtained a similar number of food pellets during the food training task (Figure 2C), indicating that group differences in self-administration behavior are likely not due to differences in associative learning of the operant task. To assess whether the increase in consumption of a food reward extended to other rewarding domains, such as drug rewards, we tested a second cohort of rats in a long-access FR1 cocaine self-administration paradigm for 6 h/day for 10 days. Similarly to what we observed with sucrose self-administration, hPMCA rats obtained more cocaine infusions across test days (Figure 2D). However, there was no difference in the number of active lever presses or inactive lever presses (Figure 2E) and no difference in food training behavior (Figure 2F) between groups. Together, these data demonstrate that blunting calcium signaling in NAc core astrocytes increases the consumption of food and drug rewards, without affecting operant task learning.

**Figure 2.**
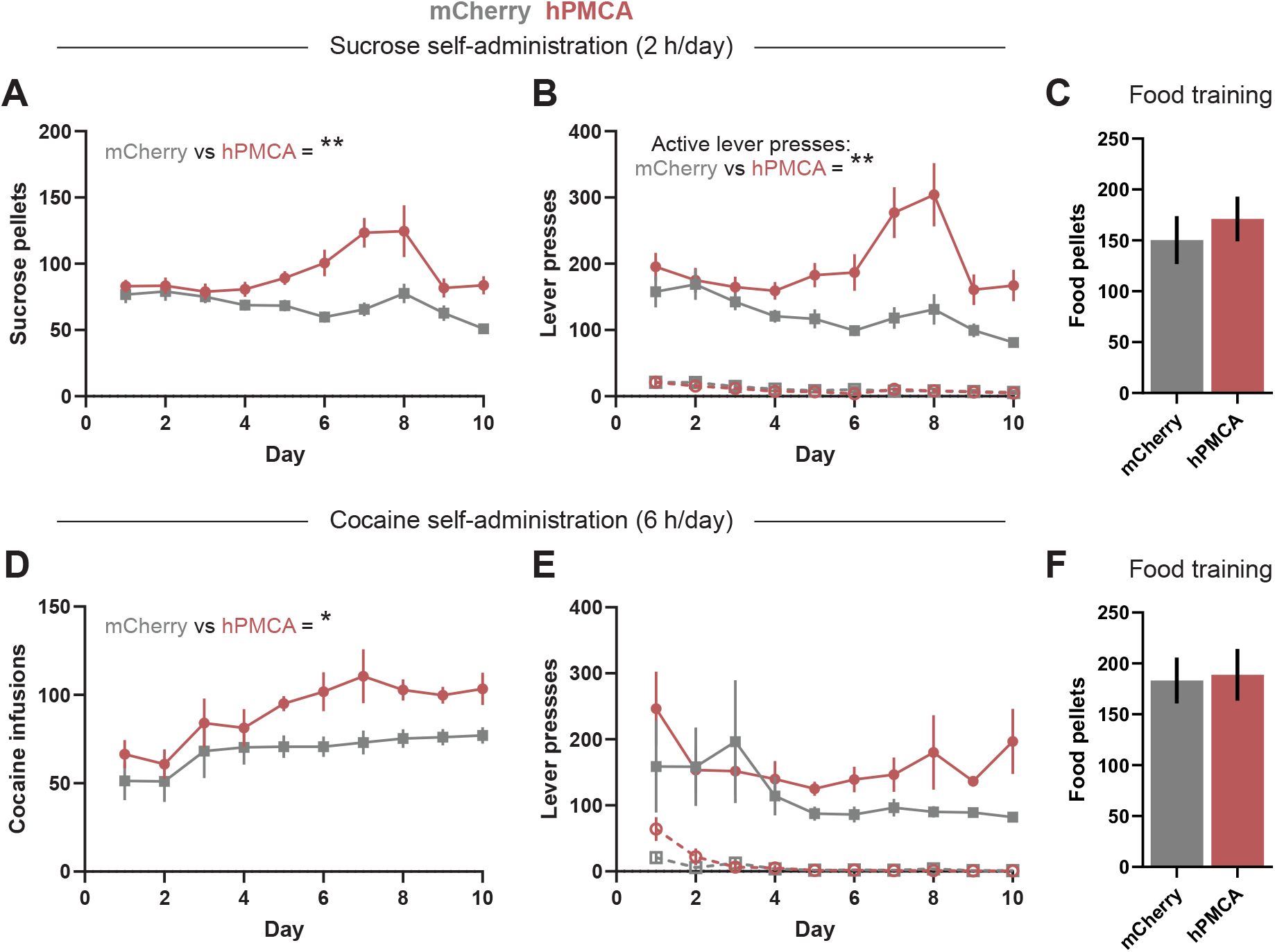
Expression of hPMCA in NAc core astrocytes alters sucrose and cocaine self-administration behavior. **A-C)** Sucrose self-administration: **(A)** hPMCA-expressing rats obtained more sucrose pellet rewards (two-way ANOVA main effect of group F(1, 16) = 15.36; p = 0.0012), and had **(B)** greater active lever pressing (solid lines) across self-administration test days (two-way ANOVA main effect of group F(1, 16) = 13.49; p = 0.0021), but had no change in inactive lever pressing (dashed lines) across test days. **(C)** Groups did not differ in the number of food pellets obtained during food training prior to self-administration. **D-F)** Cocaine self-administration: **(D)** hPMCA-expressing rats obtained more cocaine infusions across test days (two-way ANOVA main effect of group F(1, 12) = 7.253; p = 0.0196), but had no change in either **(E)** active lever pressing (solid lines) or inactive lever pressing (dashed lines). **(F)** Groups did not differ in the number of food pellets obtained during food training prior to self-administration. n = 9 rats/group in A-C; n = 7 rats/group in D-F. Points and bars represent the mean ± SEM. NAc, nucleus accumbens. *p<0.05, **p<0.01

### Expression of hPMCA in nucleus accumbens core astrocytes alters economic demand for sucrose and cocaine

To determine if the observed increase in consumption of sucrose and cocaine was attributable to changes in reward valuation, we tested a separate cohort of rats on a behavioral economics task that measures reward consumption across an increasing price schedule for sucrose or cocaine to quantify parameters related economic demand. When sucrose was the reward, hPMCA rats exhibited lower α (i.e. reduced demand elasticity) (Figure 3A, 3B), indicating that hPMCA rats were more motivated to maintain their preferred level of sucrose intake at higher ‘prices’. Consistent with this, hPMCA rats displayed a higher P_Max_ (maximum price paid for reward) (Figure 3C) and higher essential value for sucrose (i.e. stronger reinforcer strength) (Figure 3D), compared to controls. Q_0_ (theoretical consumption at zero cost) was similar between groups (Figure 3E). When cocaine was the reward, we observed similar changes in behavior: hPMCA rats exhibited lower α (Figure 3F, 3G), a near-significant increase in P_Max_ (p = 0.0519; Figure 3H), increased essential value (Figure 3I) increased Q_0_ (Figure 3J), . These collective changes in behavior indicate that reducing calcium signaling in NAc core astrocytes increases several measures of reward valuation for both sucrose and cocaine.

**Figure 3.**
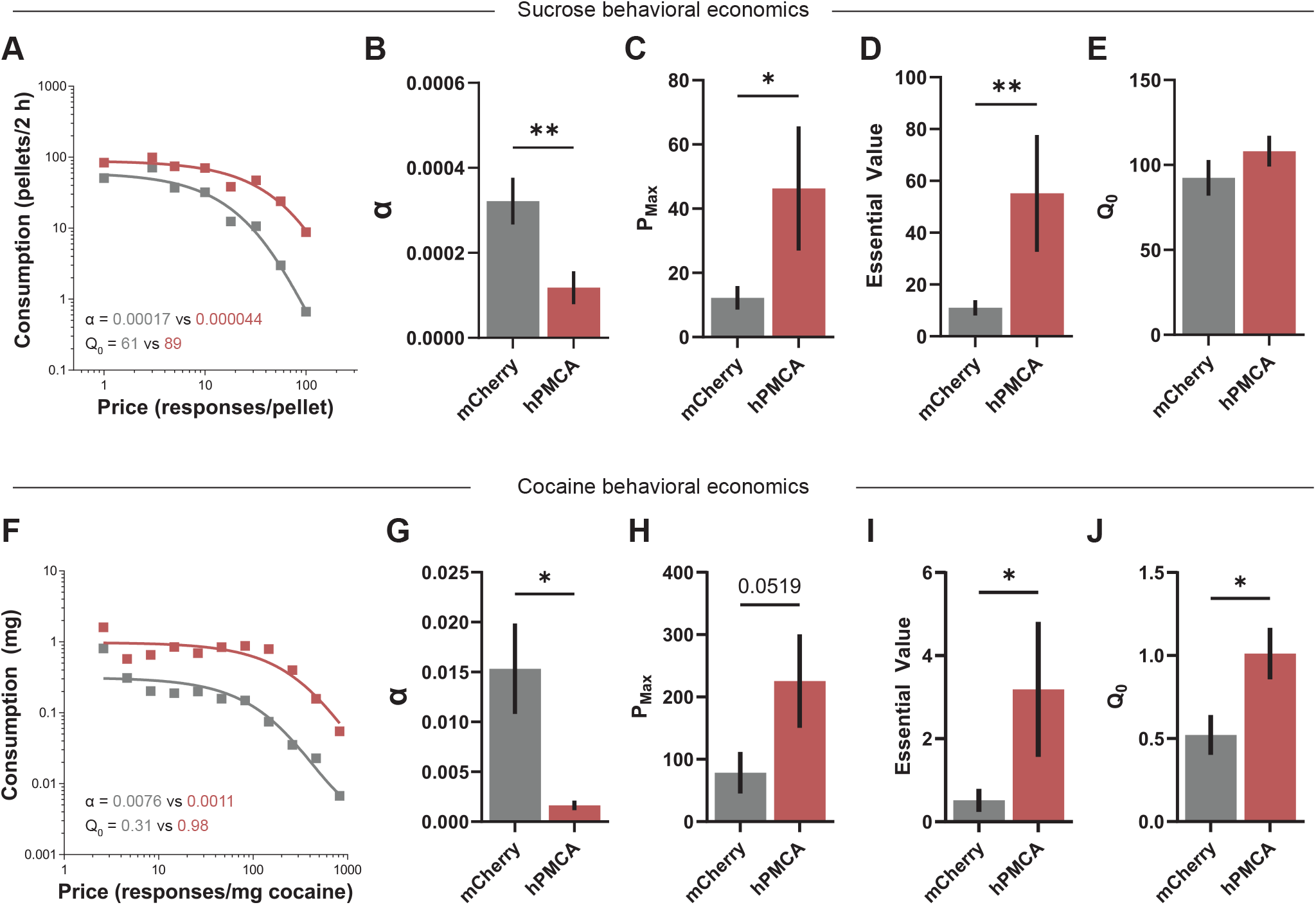
Expression of hPMCA in NAc core astrocytes alters behavioral economic parameters for both sucrose and cocaine. **A-C)** Sucrose behavioral economics: **A)** Representative group average exponential demand curves for sucrose in hPMCA-(red) and mCherry-expressing (grey) rats. **B)** hPMCA-expressing rats had reduced α indicating lower demand elasticity (increased motivation) for sucrose (t test t(16) = 3.033; p = 0.0079), **(C)** no change in Q_0_ (consumption extrapolated to zero cost), **(D)** increased P_max_ (maximum effort to obtain a sucrose pellet) (Mann-Whitney U = 17; p = 0.04), and **(E)** increased essential value (a measure of reinforcing efficacy of a reward) (Mann-Whitney U = 11.5; p = 0.0082). **F-J)** Cocaine behavioral economics: **F)** Representative group average exponential demand curves for cocaine in hPMCA-(red) and mCherry-expressing (grey) rats. **G)** hPMCA-expressing rats had reduced α indicating lower demand elasticity (increased motivation) for cocaine (t test t(9) = 2.722; p = 0.0235), **(H)** increased Q_0_ (consumption extrapolated to zero cost) (t test t(9) = 2.540; p = 0.0317), **(I)** trending increase in P_max_ (maximum effort to obtain a cocaine infusion) (Mann-Whitney U = 4; p = 0.0519), and **(J)** increased essential value (a measure of reinforcing efficacy of a reward) (Mann-Whitney U = 2; 0.0173). n = 9 rats/group in A-E; n = 5-6 rats/group in F-J. Bars represent the mean ± SEM. NAc, nucleus accumbens. *p<0.05, **p<0.01

### Expression of hPMCA in nucleus accumbens core astrocytes does not impact behavioral sensitization to cocaine

Given that hPMCA expression altered consumption and economic demand for cocaine, and astrocytes are critical to the long-term changes in NAc function following drug self-administration (Wang et al., 2022), we sought to determine whether other drug-related behaviors might also be affected. To this end, we assessed behavioral sensitization of locomotor activity following repeated exposure to cocaine. There was no difference in the distance traveled during the baseline period between hPMCA rats and mCherry controls, either pre- or post-sensitization (Figure 4A-C), and the total distance travelled, and peak locomotor activity post-cocaine injection were similar between groups at both time points (Figure 4D, 4E). Moreover, both hPMCA rats and mCherry controls exhibited increased locomotor activity in response to cocaine post-sensitization (Figure 4A-B, 4D-E), indicating that both groups experienced robust behavioral sensitization of equal magnitude.

**Figure 4.**
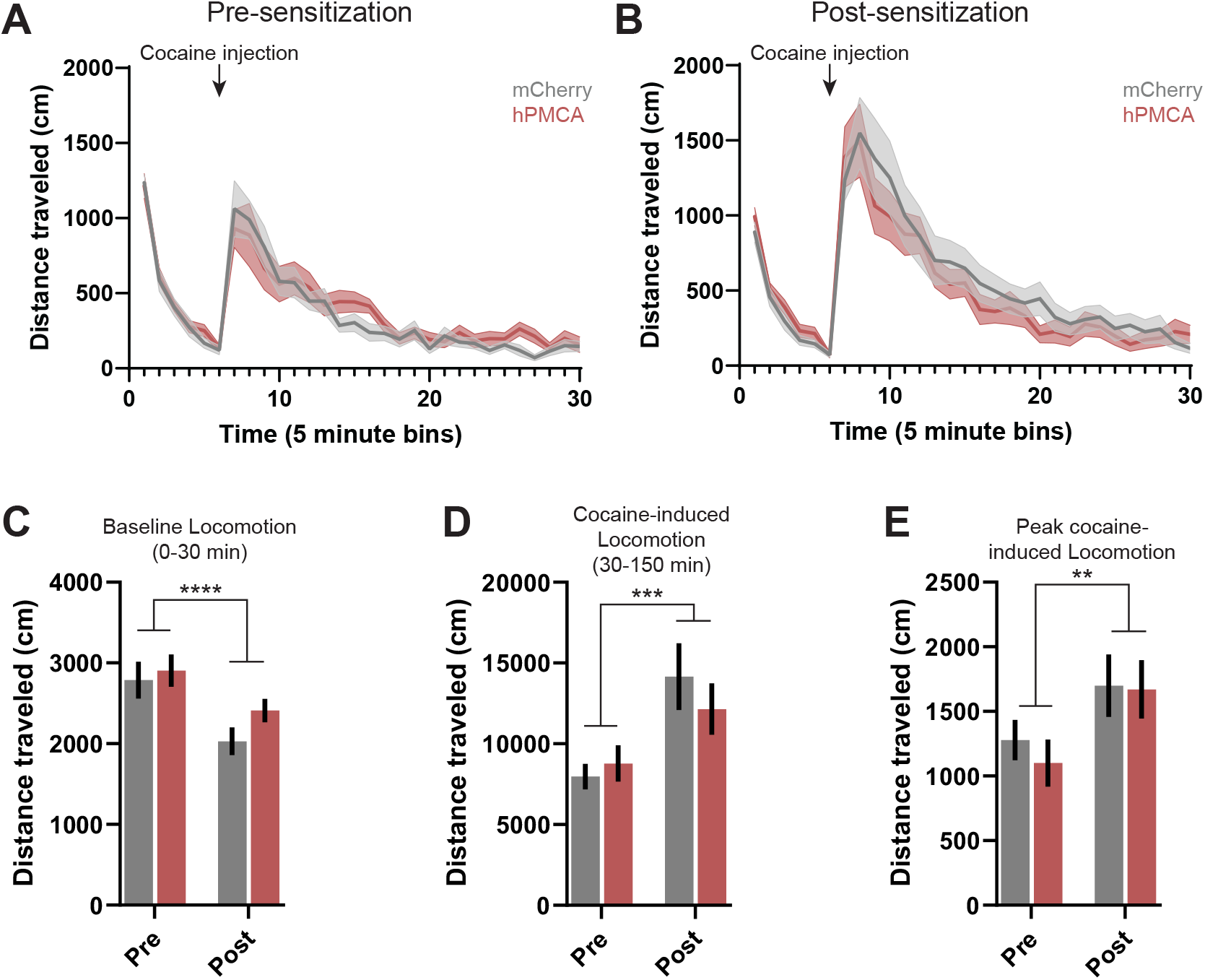
Expression of hPMCA in NAc core astrocytes does not impact behavioral sensitization to cocaine. **A-B)** Total distance traveled across the duration of the locomotor test **(A)** pre-sensitization and **(B)** post-sensitization. Arrow indicates the time at which rats were injected with cocaine (30 min into the test). **C)** The distance traveled was lower in hPMCA- and mCherry-expressing rats post-sensitization during the first 30 min baseline phase (two-way RM ANOVA main effect of test phase F(1, 24) = 25.21; p < 0.0001); however, both hPMCA- and mCherry-rats expressing rats **(D)** increased total distance traveled in response to cocaine injection post-sensitization (two-way RM ANOVA main effect of test phase F(1, 24) = 21.44; p = 0.0001) and **(E)** increased the maximum distance traveled in a single time bin in response to cocaine injection post-sensitization (two-way RM ANOVA main effect of test phase F(1, 24) = 9.405; p = 0.0053). n = 13 rats/group. Lines and bars represent the mean ± SEM. NAc, nucleus accumbens. **p<0.01, ***p<0.001, ***p<0.0001.

## DISCUSSION

Astrocytes are increasingly recognized as active regulators of neural circuit function and complex behaviors (Dallérac et al., 2018; Nagai et al., 2021; Lyon & Allen, 2022). Within the NAc, astrocyte dysfunction has been implicated in addiction-like behaviors such as drug seeking and relapse vulnerability, suggesting an underlying role in reward processing (Baker et al., 2003; Scofield et al., 2016; Kruyer et al., 2019; Siemsen et al., 2019; Kruyer & Scofield, 2021; Harder et al., 2024; Testen et al., 2025). Here, we provide evidence that astrocyte calcium signaling in the NAc core plays a central role in regulating reward consumption and valuation. Using a viral construct to express hPMCA and specifically attenuate astrocyte calcium dynamics, we found that reduced astrocyte calcium signaling led to greater self-administration of both sucrose and cocaine, along with increased behavioral economic measures of reward valuation and motivation. Importantly, altering astrocyte calcium signaling did not affect cocaine-induced locomotor sensitization, suggesting a dissociation between astrocyte function in reward processing and the neural plasticity underlying other drug-related behaviors. Together, these results highlight astrocytes within the NAc core as critical to maintaining a homeostatic “set point” for reward consumption and valuation of both natural and drug reinforcers.

Our results demonstrate that attenuating astrocyte calcium signaling increases motivated responding for reward. hPMCA rats pressed more on the active lever for both sucrose and cocaine during self-administration sessions, yet showed no differences in inactive lever pressing, indicating intact reward learning and discrimination. Furthermore, hPMCA expressing did not alter baseline locomotor activity or cocaine-induced sensitization, ruling out generalized hyperactivity as an explanation for increased active lever responding. Together, our findings suggest that NAc core astrocyte function, through intracellular calcium signaling, normally constrains reward valuation rather than influencing general motor output.

Mechanistically, reducing astrocyte calcium signaling likely impaired their ability to regulate local neural activity (Scofield & Kalivas, 2014; Corkrum & Araque, 2021). Within the NAc, neurotransmitters—such as glutamate and dopamine among others—initiate astrocyte intracellular calcium signaling cascades that lead to the release of factors that can reciprocally regulate synaptic activity and plasticity, and neural excitability (Cornell-Bell et al., 1990; Perea et al., 2009; Mariotti et al., 2016; Dallérac et al., 2018; Corkrum et al., 2020; Goenaga et al., 2023). By reducing astrocyte calcium in the present study, we likely impaired these regulatory processes resulting in an increased excitatory drive within NAc core circuits and, in turn, an increase in the valuation of rewards. This is consistent with other findings that show disrupting astrocyte calcium dynamics via gap-junction hemichannel blocker increases the motivation and consumption of ethanol following abstinence (Bull et al., 2014), and conversely, activating astrocyte calcium signaling via hM3D-Gq DREADD stimulation reduces motivation and consumption of ethanol (Bull et al., 2014) and decreases cue-induced reinstatement of cocaine seeking (Scofield et al., 2015). Together, these studies support the view that astrocytes act as a “gain control” for valuation across a range of reinforcers.

The observed increases in sucrose and cocaine consumption under low-effort reinforcement conditions in hPMCA-expressing rats parallels the heightened drug seeking and astrocytic dysfunction commonly reported in models of substance use disorders and relapse (Kruyer & Scofield, 2021), suggesting that impaired astrocytic regulation in the NAc may broadly enhance motivational drive toward rewards. While our experiments focused on sucrose and cocaine, natural reinforcers, such as food, water, sugar, sex, and social interactions share partially overlapping neural substrates with drugs of abuse in the NAc (Pitchers et al., 2010; Cameron & Carelli, 2012; Dölen et al., 2013; Bobadilla et al., 2017; Bobadilla et al., 2020; Tan et al., 2024), making it likely that broadly impairing astrocyte function would yield similar outcomes for a variety of reinforcers. Thus, astrocyte dysfunction may underlie vulnerability not only to addictive drugs, but also to conditions such as compulsive overeating, obesity, or the pathological pursuit of natural rewards in psychiatric disorders (Olsen, 2011; Zald & Treadway, 2018; Grimm, 2020). Future research is needed to determine whether reinforcing stimuli, such as social encounters, high-fat food, or other classes of drugs engage similar astrocytic mechanisms.

Interestingly, we found that reducing NAc astrocyte calcium did not alter cocaine-induced locomotor sensitization. Sensitization to cocaine depends largely on dopaminergic adaptations, structural plasticity of neurons, and AMPA receptor trafficking within the NAc (Pierce et al., 1996; Thomas et al., 2001; Li et al., 2004; Boudreau & Wolf, 2005; Scofield et al., 2016; Dietz et al., 2012). Although astrocytes influence neuronal structural plasticity and changes in glutamate homeostasis following repeated cocaine exposure, (Baker et al., 2003; Kalivas, 2009; Reissner et al., 2015; O’Donovan et al., 2021; Wang et al., 2021; Wang et al., 2022; Mongrédien et al., 2025), our findings suggest that mechanisms of sensitization may occur independently from astrocyte calcium-dependent functions or perhaps are less sensitive to astrocytic dysfunction. This dissociation highlights the role of astrocytes in regulating drug consumption and valuation without globally altering all drug-related plasticity.

Although our findings demonstrate a key role for astrocytic calcium signaling in reward-related behavior, they also raise important questions about cellular and circuit-level processes through which these effects emerge. The hPMCA construct provides a constitutive suppression of intracellular calcium, and as such, does not reveal how temporal dynamics or sub-cellular microdomain calcium signaling contribute to behavior (Bazargani & Attwell, 2016). Because astrocytes form extensive networks that influence neural activity across both local synapses and broader circuit populations, it is unclear how broadly suppressing calcium signaling in astrocytes affects specific cell types and pathways within the NAc. Reward encoding in the NAc relies on the integration of signals in a cell-type and circuit-specific manner, raising the possibility that diminished astrocytic calcium signaling may differentially impact D1-versus D2-expressing medium spiny neurons or selectively modulate input pathways from regions such as the prefrontal cortex, amygdala, or ventral tegmental area (Floresco, 2015). Clarifying these relationships will be critical for understanding how astrocytic signaling shapes reward computation at the circuit level.

## Conclusion

In conclusion, we show that reducing astrocyte calcium signaling in the NAc core increased consumption and economic valuation of sucrose and cocaine, while leaving cocaine-induced locomotor sensitization intact. These results demonstrate that astrocytic function, through intracellular calcium signaling, normally serves to constrain reward valuation and motivational drive. Together, this work adds to a growing body of evidence highlighting astrocytes as critical regulators of natural and drug reward-related behaviors. A more complete understanding of astrocytic function in reward may guide novel therapeutic strategies for substance use disorders and other disorders of motivation.

## DATA AVAILABILITY STATEMENT

The data that support the findings of this study are available from the corresponding author upon reasonable request.

## AUTHOR CONTRIBUTIONS

Conceptualization: JVR, KJR; Methodology: JVR, UG, MHJ, KJR; Formal Analysis: JVR; Investigation: JVR, GAL; Writing-Original Draft: JVR, KJR; Writing-Review and Editing: JVR, UG, MHJ, KJR; Supervision: JVR, KJR; Funding Acquisition: KJR.

## REFERENCES

Baker DA, McFarland K, Lake RW, Shen H, Tang X-C, Toda S, Kalivas PW. Neuroadaptations in cystine-glutamate exchange underlie cocaine relapse. Nat Neurosci. 2003; 6:743–749.

Bazargani N, Attwell D. Astrocyte calcium signaling: the third wave. Nature Neuroscience. 2016; 19:182–189.

Berendse HW, Galis-de Graaf Y, Groenewegen HJ. Topographical organization and relationship with ventral striatal compartments of prefrontal corticostriatal projections in the rat. J Comp Neurol. 1992; 316:314–347.

Bobadilla AC, Garcia-Keller C, Heinsbroek JA, Scofield MD, Chareunsouk V, Monforton C, Kalivas PW. Accumbens mechanisms for cued sucrose seeking. Neuropsychopharmacology. 2017; 43:2377–2386.

Bobadilla AC, Dereschewitz E, Vaccaro L, Heinsbroek JA, Scofield MD, Kalivas PW. Cocaine and sucrose rewards recruit different seeking ensembles in the nucleus accumbens core. Molecular Psychiatry. 2020; 25:3150–3163.

Boudreau AC, Wolf ME. Behavioral sensitization to cocaine is associated with increased AMPA receptor surface expression in the nucleus accumbens. J Neurosci. 2005; 25:9144–9151.

Britt JP, Benaliouad F, McDevitt RA, Stuber GD, Wise RA, Bonci A. Synaptic and behavioral profile of multiple glutamatergic inputs to the nucleus accumbens. Neuron. 2012; 76:790–803.

Bull C, Freitas KCC, Zou S, Poland RS, Syed WA, Urban DJ, Minter SC, Shelton KL, Hauser KF, Negus SS, Knapp PE, Bowers MS. Rat nucleus accumbens core astrocytes modulate reward and the motivation to self-administer ethanol after abstinence. Neuropsychopharmacology. 2014; 39:2835–2845.

Calipari ES, Bagot RC, Purushothaman I, Davidson TJ, Yorgason JT, Pena CJ, Walker DM, Pirpinias ST, Guise KG, Ramakrishnan C, Deisseroth K, Nestler EJ. In vivo imaging identifies temporal signature of D1 and D2 medium spiny neurons in cocaine reward. Proc Natl Acad Sci U S A. 2016; 113:2726–2731.

Cameron CM, Carelli RM. Cocaine abstinence alters nucleus accumbens firing dynamics during goaldirected behaviors for cocaine and sucrose. European Journal of Neuroscience. 2012; 35:940–051.

Corkrum M, Covelo A, Lines J, Bellocchio L, Pisansky M, Loke K, Quintana R, Rothwell PE, Lujan R, Marsicano G, Martin ED, Thomas MJ, Kofuji P, Araque A. Dopamine-evoked synaptic regulation in the nucleus accumbens requires astrocyte activity. Neuron. 2020; 105:1036-1047.e5.

Corkrum M, Araque A. Astrocyte-neuron signaling in the mesolimbic dopamine system: the hidden stars of dopamine signaling. Neuropsychopharmacology. 2021; 46:1864–1872.

Cornell-Bell AH, Finkbeiner SM, Cooper MS, Smith SJ. Glutamate induces calcium waves in cultured astrocytes: long-range glial signaling. Science. 1990; 247:470–473.

Dallérac G, Zapata J, Rouach N. Versatile control of synaptic circuits by astrocytes: where, when and how? Nat Rev Neurosci. 2018; 19:729–743.

Dietz DM, Sun H, Lobo MK, Cahill ME, Chadwick B, Gao V, Koo JW, Mazei-Robison MS, Dias C, Maze I, Damez-Werno D, Dietz KC, Scobie KN, Ferguson D, Christoffel D, Ohnishi Y, Hodes GE, Zheng Y, Neve RL, Hahn KM, Russo SJ, Nestler EJ. Rac1 is essential in cocaine-induced structural plasticity of nucleus accumbens neurons. Nat Neurosci. 2012; 15:891–896.

Dölen G, Darvishzadeh A, Huang KW, Malenka RC. Social reward requires coordinated activity of accumbens oxytocin and 5HT. Nature. 2013; 501:179–184.

Erickson EK, DaCosta AJ, Mason SC, Blednov YA, Mayfield RD, Harris RA. Cortical astrocytes regulate ethanol consumption and intoxication in mice. Neuropsychopharmacology. 2020; 46:500–508.

Floresco SB. The nucleus accumbens: an interface between cognition, emotion, and action. Annu Rev Psychol. 2015; 66:25–52.

Goenaga J, Araque A, Kofuji P, Herrera Moro Chao D. Calcium signaling in astrocytes and gliotransmitter release. Front Synaptic Neurosci. 2023; 15:1138577.

Grimm JW. Incubation of food craving in rats: a review. J Exp Anal Behav. 2020; 113:37–47.

Harder EV, Franklin JP, VanRyzin JW, Reissner KJ. Astrocyte-neuron interactions in substance use disorders. In: Blanco-Suarez E, Farhy-Tselnicker I, editors. Astrocyte-neuron interactions in health and disease. Advances in Neurobiology, vol 39. Springer, Cham; 2024. p. 165–191.

Heinsbroek JA, Giannotti G, Mandel MR, Josey M, Aston-Jones G, James MH, Peters J. A common limiter for opioid choice and relapse identified in a rodent addiction model. Nat Commun. 2021; 12:4788.

Hursh SR, Silberberg A. Economic demand and essential value. Psychol Rev. 2008; 115:186–198.

Kalivas PW, Duffy P. Effect of acute and daily cocaine treatment on extracellular dopamine in the nucleus accumbens. Synapse. 1990; 5:48–58.

Kalivas PW. The glutamate homeostasis hypothesis of addiction. Nat Rev Neurosci. 2009; 10:561–572.

Kastner-Blasczyk AR, Hester SW, Reasons SE, Scofield MD, Woodward JJ. Effect of an astrocyte calcium exporter on orbitofrontal cortex neuron excitability, astrocyte-synaptic interaction, and alcohol consumption. Neuropharmacology. 2025; 269:110365.

Kruyer A, Scofield MD, Wood D, Reissner KJ, Kalivas PW. Heroin cue-evoked astrocytic structural plasticity at nucleus accumbens synapses inhibits heroin seeking. Biol Psychiatry. 2019; 86:811–819.

Kruyer A, Scofield MD. Astrocytes in addictive disorders. Adv Neurobiol. 2021; 26:231–254.

Kutlu MG, Zachry JE, Melugin PR, Cajigas SA, Chevee MF, Kelly SJ, Kutlu B, Tian L, Siciliano CA, Calipari ES. Dopamine release in the nucleus accumbens core signals perceived saliency. Current Biology. 2021; 31:4748-4761.e8.

Li Y, Acerbo MJ, Robinson TE. The induction of behavioural sensitization is associated with cocaine-induced structural plasticity in the core (but not shell) of the nucleus accumbens. Eur J Neurosci. 2004; 20:1647–1654.

Li Z, Chen Z, Fan G, Li A, Yuan J, Xu T. Cell-type-specific afferent innervation of the nucleus accumbens core and shell. Front Neuroanat. 2018; 12:84.

Lobo MK, Covington 3rd HE, Chaudhury D, Friedman AK, Sun HS, Damez-Werno D, Dietz DM, Zaman S, Koo JW, Kennedy PJ, Mouzon E, Mogri M, Neve RL, Deisseroth K, Han M-H, Nestler EJ. Cell type-specific loss of BDNF signaling mimics optogenetic control of cocaine reward. Science. 2010; 330:385–390.

Lyon KA, Allen NJ. From synapses to circuits, astrocytes regulate behavior. Front Neural Circuits. 2022; 15:786293.

Mariotti L, Losi G, Sessolo M, Marcon I, Carmignoto G. The inhibitory neurotransmitter GABA evokes long-lasting Ca(2+) oscillations in cortical astrocytes. Glia. 2016; 64:363–373.

Mogenson GJ, Jones DL, Yim CY. From motivation to action: functional interface between the limic system and the motor system. Prog Neurbiol. 1980; 14:69–97.

Mongrédien R, Anesio A, Fernandes GJD, Eagle AL, Maldera S, Pham C, Robert S, Bezerra F, Vilette A, Bianchi PC, Franco C, Louis F, Gruszczynski C, Niépon M-L, Betancur C, Erdozian AM, Robison AJ, Boucard AA, Cruz FC, Li D, Heck N, Gautron S, Vialou V. Astrocytes control cocaine-induced synaptic plasticity and reward through the matricellular protein Hevin. Biol Psychiatry. 2025; 3223:01072–8.

Nagai J, Yu X, Papouin T, Cheong E, Freeman MR, Monk KR, Hastings MH, Haydon PG, Rowitch D, Shaham S, Khakh BS. Behaviorally consequential astrocytic regulation of neural circuits. Neuron. 2021; 109:576–596.

O’Donovan B, Neugornet A, Neogi R, Xia M, Ortinski PI. Cocaine experience induces functional adaptations in astrocytes: implications for synaptic plasticity in the nucleus accumbens shell. Addict Biol. 2021; 26:e13042.

Olsen CM. Natural rewards, neuroplasticity, and non-drug addictions. Neuropharmacology. 2011; 61:1109–1122.

Papouin T, Dunphy J, Tolman M, Foley JC, Haydon PG. Astrocytic control of synaptic function. Philos Trans R Soc Lond B Biol Sci. 2017; 273:20160154.

Perea G, Navarrete M, Araque A. Tripartite synapses: astrocytes process and control synaptic information. Trends Neurosci. 2009; 32:421–431.

Pierce RC, Bell K, Duffy P, Kalivas PW. Repeated cocaine augments excitatory amino acid transmission in the nucleus accumbens only in rats having developed behavioral sensitization. J Neurosci. 1996. 16:1550–1560.

Pitchers KK, Balfour ME, Lehman MN, Richtand NM, Yu L, Coolen LM. Neuroplasticity in the mesolimbic system induced by natural reward and subsequent reward abstinence. Biol Psychiatry. 2010; 67:872–879.

Reissner KJ, Gipson CD, Tran PK, Knackstedt LA, Scofield MD, Kalivas PW. Glutamate transporter GLT-1 mediates N-acetylcysteine inhibition of cocaine reinstatement. Addict Biol. 2015; 20:316–323.

Scofield MD, Kalivas PW. Astrocytic dysfunction and addiction: consequences of impaired glutamate homeostasis. Neuroscientist. 2014; 20:610–22.

Scofield MD, Heinsbroek JA, Gipson CD, Kupchik YM, Spencer S, Smith ACW, Roberts-Wolfe D, Kalivas PW. The nucleus accumbens: Mechanisms of addiction across drug classes reflect the importance of glutamate homeostasis. Pharmacol Rev. 2016; 68:816–871.

Scofield MD, Li H, Siemsen BM, Healey KL, Tran PK, Woronoff N, Boger HA, Kalivas PW, Reissner KJ. Cocaine self-administration and extinction leads to reduced glial fibrillary acidic protein expression and morphometric features of astrocytes in the nucleus accumbens core. Biol Psychiatry. 2016; 80:207–215.

Siemsen BM, Reichel CM, Leong KC, Garcia-Keller C, Gipson CD, Spencer S, McFaddin JA, Hooker KN, Kalivas PW, Scofield MD. Effects of methamphetamine self-administration and extinction on astrocyte structure and function in the nucleus accumbens core. Neuroscience. 2019; 406:528–541.

Soares-Cunha C, Coimbra B, David-Pereira A, Borges S, Pinto L, Costa P, Sousa N, Rodrigues AJ. Activation of D2 dopamine receptor-expressing neurons in the nucleus accumbens increases motivation. Nat Commun. 2016; 7:11829.

Stuber GD, Sparta DR, Stamatakis AM, van Leeuwen WA, Hardjoprajitno JE, Cho S, Tye KM, Kempadoo KA, Zhang F, Deisseroth K, Bonci A. Excitatory transmission from the amygdala to nucleus accumbens facilitates reward seeking. Nature. 2011; 475:377–380.

Tan B, Browne CJ, Nobauer T, Vaziri A, Friedman JM, Nestler EJ. Drugs of abuse hijack a mesolimbic pathway that processes homeostatic need. Science. 2024; 384:eadk6742.

Tavakoli NS, Malone SG, Anderson TL, Neeley RE, Asadipooya A, Bardo MT, Otinski PI. Astrocyte Ca2+ in the dorsal striatum suppresses neuronal activity to oppose cue-induced reinstatement of cocaine seeking. Front Cell Neurosci. 2024; 18:1347491.

Testen A, VanRyzin JW, Bellinger TJ, Kim R, Wang H, Gastinger MJ, Witt EA, Franklin JP, Vecchiarelli HA, Picard K, Scofield MD, Tremblay M-È, Reissner KJ. Abstinence from cocaine self-administration promotes microglial pruning of astrocytes, which drives cocaine-seeking behavior. Cell Reports. 2025; 44:116137.

Thomas MJ, Beurrier C, Bonci A, Malenka RC. Long-term depression in the nucleus accumbens: a neural correlate of behavioral sensitization to cocaine. Nat Neurosci. 2001; 4:1217–1223.

Wang J, Li K-L, Shukla A, Beroun A, Ishikawa M, Huang X, Wang Y, Wang YQ, Yang Y, Bastola ND, Huang HH, Kramer LE, Chao T, Huang YH, Sesack SR, Nestler EJ, Schluter OM, Dong Y. Cocaine triggers astrocyte-mediated synaptogenesis. Biol Psychiatry. 2021; 89:386–397.

Wang J, Holt LM, Huang HH, Sesack SR, Nestler EJ, Dong Y. Astrocytes in cocaine addiction and beyond. Molecular Psychiatry. 2022; 27:652–668.

Yu X, Taylor AMW, Nagai J, Golshani P, Evans CJ, Coppola G, Khakh BS. Reducing astrocyte calcium signaling in vivo alters striatal microcircuits and causes repetitive behavior. Neuron. 2018; 99:1170–1187e9.

Zald DH, Treadway M. Reward processing, neuroeconomics, and psychopathology. Annu Rev Clin Psychol. 2017; 13:471–495.

